# A Structural Design Principle for Temperature Robustness in Biomolecular Circuits

**DOI:** 10.64898/2026.08.14.744825

**Authors:** Gunjan Chorasiya, Shaunak Sen

## Abstract

The dominant paradigm for temperature robustness in biomolecular circuits is for the parameters to be tuned to have matching temperature dependencies so that their overall effect cancels out. This contrasts with the robustness due to circuit structure, typically operative in circuits where robustness to a single input parameter is desired. The importance of the circuit structure in temperature robustness is generally unclear. We addressed this issue in a benchmark negative feedback circuit using a combination of theoretical modelling and experimental measurements. We found that the response to a temperature perturbation in a model of negative feedback was qualitatively different from the response in a model without feedback. We experimentally measured the response of the negative feedback circuit to a temperature perturbation and found that it was smaller than that of the circuit without feedback, in line with the theoretical finding. We confirmed this theoretical prediction experimentally. The initial response of the negative feedback circuit, paradoxically, was larger than the circuit without feedback. The resolution of this paradox was in accounting for the faster dynamics in the negative feedback circuit. These results show a simple design principle of temperature robustness that can operate in a widespread circuit motif and may also apply to other perturbations which, like temperature, affect multiple parameters simultaneously.

## I. Introduction

Uncovering the principles of temperature compensation is a fundamental problem in biology. The classical example of temperature compensation, or robustness, is the time period of circadian oscillations that is relatively unchanged over a wide range of temperatures [1], [2]. The underlying principle is understood to be the “matching” of the temperature dependencies of different parameters so that the overall effect of a temperature change cancels out at the functional output [3], [4], [5], [6], [7], [8]. There have been attempts to investigate structural, adaptive sources of temperature robustness that do not depend on parameters [9], [10] (see also [11], [12]). Designing temperature robustness is an important specification in synthetic biology as well [13]. The parameter matching principle has been used to experimentally design temperature robustness in a synthetic biomolecular oscillator [14]. More generally, the parameter matching principle is used to facilitate temperature robustness in multiple engineering contexts (see for example [15]). In biological contexts, however, measuring exact parameter values, let alone their temperature dependence, as well as rationally designing parameters to meet a pre-specified temperature dependence can be quite challenging.

While parameter matching is the dominant paradigm for temperature robustness, it is the circuit structure, typically negative feedback and variants, that are widely held to underlie robustness in biomolecular contexts, especially when robustness to a single input perturbation is desired. In fact, negative feedback has been shown to be one of two ways, the other being feedforward, to implement exact adaptation to a single input perturbation in a class of biomolecular circuits [16]. The canonical studies of robustness in bacterial chemotaxis and the identification of an integral negative feedback control as the core mechanism driving this robustness were pivotal steps in the quantitative development of systems biology [17], [18]. An experimental demonstration of integral feedback in a cell-free environment showed robust reference tracking and disturbance rejection in synthetic gene circuits, including to temperature [19]. Robustness against a temperature perturbation is qualitatively different from the robustness to a single input signal or concentration because temperature can impact multiple parameters of a biomolecular circuit (Fig. 1(a)). Paradoxically, the parameters of the “controller” module designed to enhance the robustness may themselves vary with temperature.

**Fig. 1:**
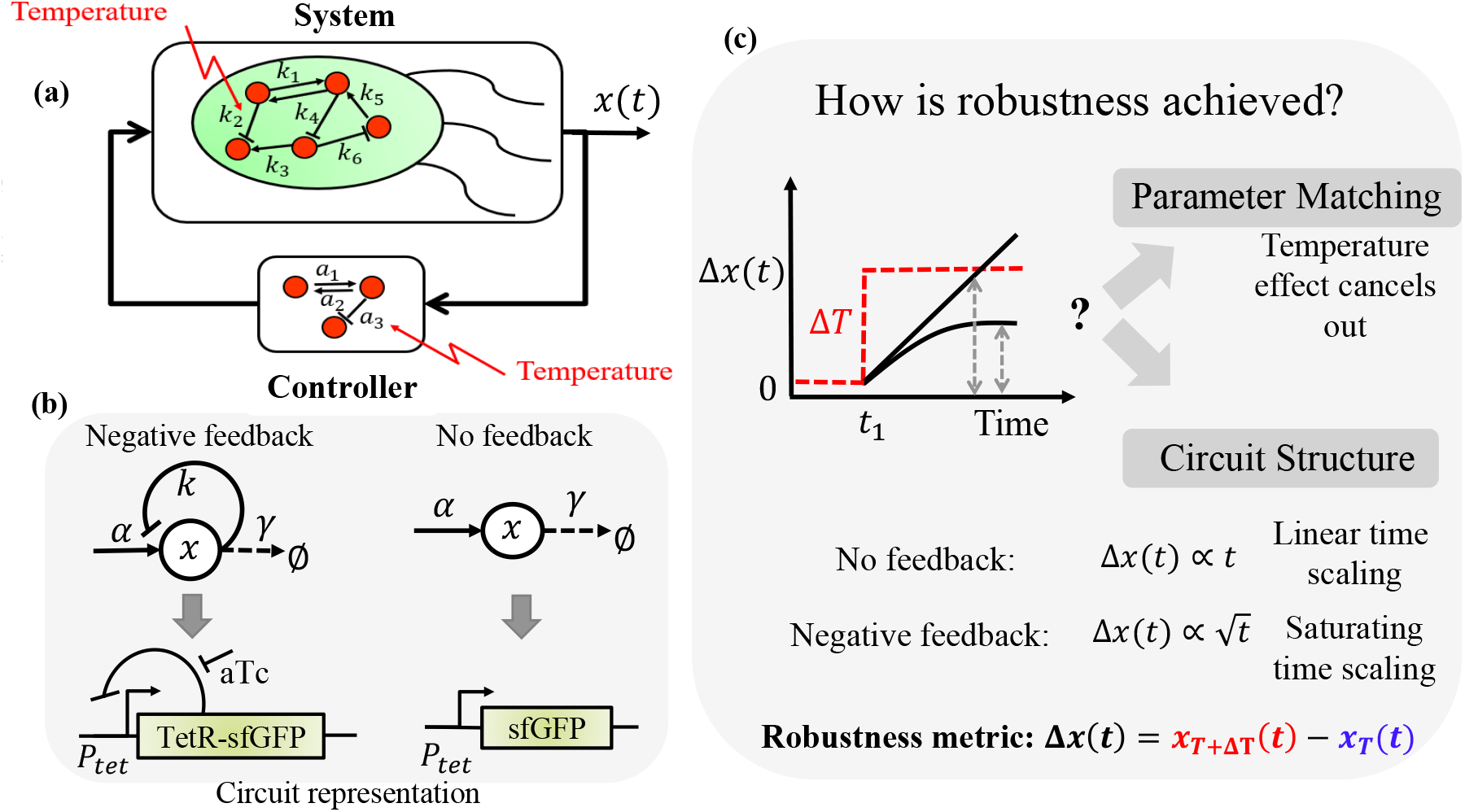
Temperature Robustness in Biomolecular Circuits. (a) Illustration of a cell as a system and controller; both are sensitive to temperature, for example through the indicated rate constants. (b) The representation of a negative feedback circuit and a circuit without feedback in computations and in experiments. (c) The overview of the mechanisms of robustness to step-change in temperature. The dotted red line denotes a step change in temperature. The symbol Δ*x*(*t*) denotes the difference in trajectories between the response when a temperature step is applied *x*_*T* +Δ*T*_ (*t*) and the nominal temperature response *x*_*T*_ (*t*). The dotted grey lines represent the robustness metric; the smaller Δ*x*(*t*) represents higher robustness.

Whether and how circuit structure can facilitate temperature robustness is relatively unclear (see also [20]). There are at least three lines of evidence that point to the existence of a design principle for temperature robustness that does not rely on exact parameter matching. One, the cell-free implementation of integral feedback exhibited robustness to temperature, even though there was no explicit design for matching temperature dependence of parameters [19]. Two, we have presented theoretical and experimental work showing that the structural nonlinearities in biomolecular circuits could enable temperature robustness in certain parameter regimes [21], [22]. Recent work on temperature dependence in biomolecular contexts also underlines the importance of nonlinear effects [23]. Three, investigations of biomolecular noise, a perturbation similar to temperature in impacting multiple circuit parameters, had shown how the steady-state distribution in a negative feedback circuit was more robust than in a circuit without feedback [24]. While parameter matching could be the predominant principle for designing temperature robustness, a structural principle based on operative parameter regimes may be a useful metaphor for analysis and design.

The main idea we used to study whether temperature robustness could originate from the circuit structure was to study the effect of a step change in temperature on the circuit response: a more robust circuit structure would show a smaller difference in the response. Our approach was a combination of experimental measurements in living *E. coli* cells and the analysis of mathematical models using theoretical methods and computer simulations. Using simple mathematical models, we predicted that a transcriptional negative feedback circuit would show a smaller difference in the response in comparison with a circuit without the negative feedback. We experimentally verified this prediction by comparing the response of a transcriptional negative feedback circuit with a circuit without feedback. As predicted, we found that the difference in the response was proportional to time in the circuit without feedback and exhibited a saturation effect as a function of time in the circuit with the feedback. Counterintuitively, we found that while the negative feedback circuit was more robust than the no feedback circuit, the initial rate of divergence was actually higher. The resolution of this paradox was in reconciling two opposing consequences of increasing negative feedback: faster dynamics that contract more. These results provided direct evidence for and a deeper understanding of how temperature robustness originates from the circuit structure.

## II. Results

### A. Mathematical Model

While negative feedback is synonymous with robustness, its efficacy in robustness to a temperature perturbation is generally unclear because the parameters implementing the negative feedback could themselves change with temperature. Therefore, we started our investigation with the canonical transcriptional negative feedback circuit (Fig. 1(b)). This negative autoregulation circuit has been identified as a transcriptional feedback motif, underlining its importance in transcriptional regulation [25]. To abstract the essence of the negative feedback, we used a simple mathematical model [26], [27]. The mathematical model consisted of a transcription factor *X* that repressed its own expression,

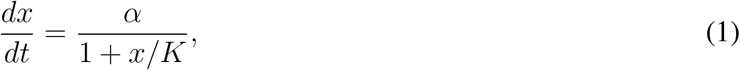

where *x* denoted the concentration of the transcription factor and its rate of change was given by a saturating Hill function with maximum production rate *α* and DNA binding constant *K*. As the concentration of *X* increased, it’s rate of production decreased, capturing the essence of negative feedback. In contrast, a circuit without feedback would have a constant rate of production,

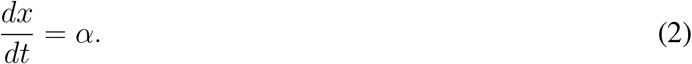

In these models, the parameters *α* and *K* could depend on temperature and the key point of our investigation is how the responses to a temperature feedback differ with and without feedback (see also [20]).

We found that the transcriptional negative feedback circuit was more robust to a step change in temperature than a circuit without feedback (Fig. 1(c)). We considered the robustness to a step change in temperature because it was a perturbation that was operationally well-defined in both theoretical and experimental contexts. We assessed the extent of robustness by the difference in the response with and without the temperature step. For the circuit without feedback,

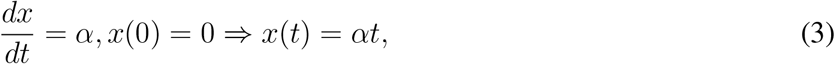

and the difference in the response was Δ*x*(*t*) = Δ*αt*, where a temperature step *T* → *T* +Δ*T* is applied at *t* = 0 and Δ*α* = *α*(*T* + Δ*T*) − *α*(*T*) is the difference in the production parameter due to the temperature step. In this case, the difference in response increased linearly with time Δ*x*(*t*) ∼ *t*. To obtain similar analytical insight for the negative feedback model, we considered the regime of strong repression *x* ≫ *K* [27]. In this regime, 1 + *x/K* ≈ *x/K* and the simplified model is,

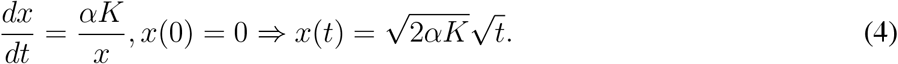

The difference in the trajectories Δ*x*(*t*) = *x*_*T* +Δ*T*_ (*t*) − *x*_*T*_ (*t*) can be calculated from 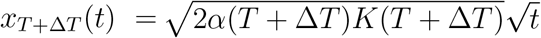, and 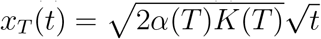. The difference is, 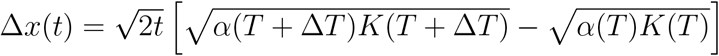.

Therefore, the difference in response follows the time-scaling 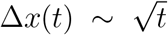, with the constant of proportionality being a function of the difference in parameters. The time scaling 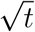 in the negative feedback grew more slowly in time than the time scaling *t* in the circuit without feedback. The different time-scalings directly follow from the nonlinearity in the negative feedback model. This makes the negative feedback more robust to a temperature step than the circuit without feedback.

The qualitative difference in the response to a temperature step between the negative feedback circuit and a circuit without feedback was a strong prediction. The overall trends persisted when the models were simulated without the above approximation (Fig. 2 (a), (b)).

**Fig. 2:**
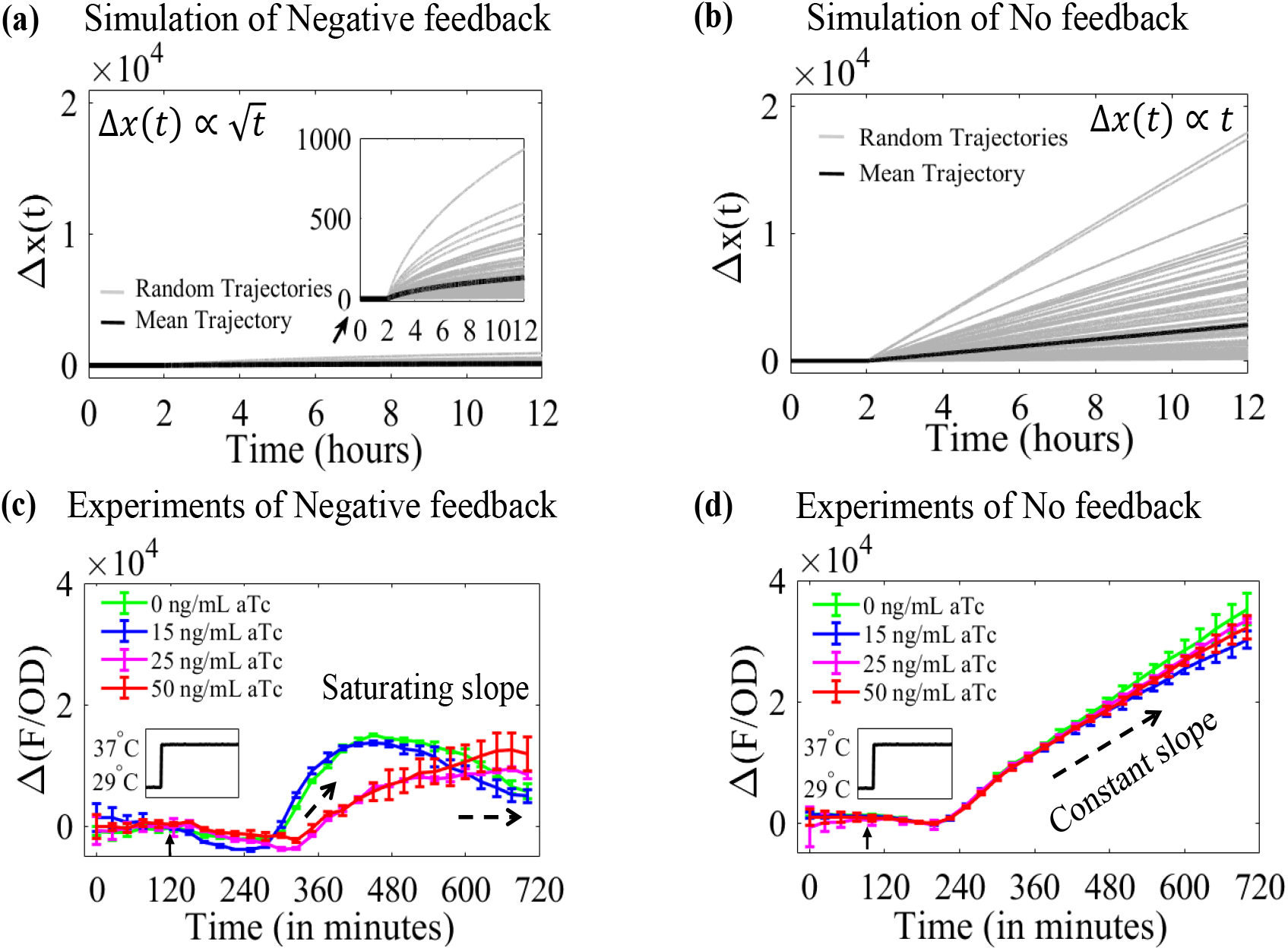
Experimental measurements of circuit responses to a temperature perturbation. The top two subfigures show the simulations of (a) the negative feedback model and (b) the no feedback model. The symbol Δ*x*(*t*) is the difference between temperature step response and control response. The grey and solid black lines in the simulations represent the number of random trajectories (*m* = 100) and their mean, respectively. The nominal parameters used were *α*_0_ = 100*nM/hr* and *K*_0_ = 10*nM*. For the simulations, these parameters were randomly sampled as 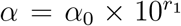 and 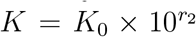, where *r*_1_ and *r*_2_ are independent uniformly distributed random variables in the interval [−1, 1]. Simulations of the differential equations were done in MATLAB using the ode45 solver. The bottom two subfigures show experimental trajectories of (c) negative feedback and (d) no feedback circuit. The small black arrow indicates the temperature step applied at the 120^*th*^ minute. The temperature step is used as a perturbation, with the step occurring at the 2^*nd*^ hour from 29°C to 37°C in both simulation and experimental results. The colors in the experimental figure represent different aTc concentrations: the red curve is at 50 *ng/mL* aTc, the magenta curve is 25 *ng/mL* aTc, the blue curve is 15 *ng/mL* aTc, and the green curve is 0 *ng/mL* aTc. The smaller deviations of the difference in trajectories in a negative feedback circuit show more robustness than a no feedback circuit.

### B. Experimental Measurements

We tested the hypothesis of temperature robustness due to the circuit structure experimentally. A transcriptional negative feedback circuit and a circuit without feedback were synthesized and the difference in their responses due to a temperature step were measured. The negative feedback circuit is the classical *P*_*tet*_ − *TetR* circuit where the Tetracycline Repressor (TetR) is fused to a Green Fluorescent Protein (GFP) and the fusion protein is placed under the control of a *P*_*tet*_ promoter [24]. This circuit allows the use of the inducer anhydrotetracycline (*aTc*) as a tuning knob for the feedback strength via the inhibition of the *TetR* activity. The circuit without feedback expressed GFP from a *P*_*tet*_ promoter. We measured the dynamics of both circuits in a microplate reader. The temperature of the microplate reader was set to 29°*C*. A temperature perturbation was applied by shifting the temperature to 37°*C* at a specified time-point. To measure the difference in response, one set of experiments was performed without the temperature step. We expected that the difference in response when the temperature changes from 29°*C* to 37°*C* would be less in the negative feedback circuit relative to that in a circuit with no feedback.

The experimentally measured difference in response to a temperature step was indeed smaller in the circuit with negative feedback compared to the circuit without feedback (Fig. 2(c), (d)). The difference in response increased almost linearly in the circuit without feedback, whereas the difference exhibited a saturating trend in the negative feedback circuit. This was in line with the modelling predictions (Fig. 1(c)). The overall trends observed persisted when minimal media was used (See Supplementary Figure 1). The raw data are shown in the Supplementary Figure 2. Temperature is a global environmental variable and could affect aspects of the measurement process such as the GFP properties. These effects, however, should be the same for both circuits and the main conclusion should hold.

These results experimentally validated the hypothesis that temperature robustness could arise from the underlying circuit structure. In the standard negative feedback configuration, the error between the desired output and the actual output is negatively fed back to the input in such a manner that the error reduces. The goal in this case is to attenuate the effect of disturbances that give rise to the error and it is assumed that the negative feedback parameters are immune to the disturbance. When temperature is the disturbance that is to be attenuated, the situation is different as the parameters of the negative feedback circuit could also depend on temperature. The robustness to temperature achieved with the negative feedback circuit points to a more general principle. This design principle for temperature robustness depends on the circuit structure and not necessarily on the cancellation of the temperature dependencies of the parameters.

## C. Discussion

Counterintuitively, the initial rate of increase of the difference in the negative feedback trajectories was larger than in the no feedback trajectories. This is clearly seen in the experimental trajectories. The initial slope of increase in the circuit without feedback is 110.15 au/ min whereas in the circuit with negative feedback this can be as high as 143.39 au/ min. In fact, even within the negative feedback circuit, the stronger feedback trajectories (low aTc) had a larger slope than the weaker feedback (high aTc) trajectories. This was counterintuitive because the overall difference in the response to the temperature step had a larger saturation effect in the negative feedback circuit than in the no feedback circuit.

We found that this apparent paradox owed its origin to two opposing tendencies of negative feedback. The first tendency is the overall contracting effect investigated in the above sections. The second tendency is the increasing response speed that makes the response to a disturbance, such as a temperature step, faster. These can be illustrated through the mathematical model

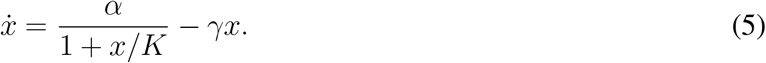

The linearisation of this model around an operating point *x*_0_ and nominal parameter set (*α, K, γ*) is

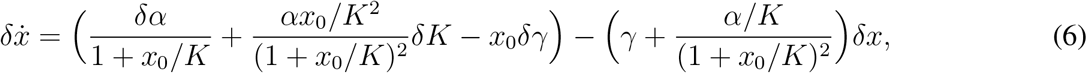

where *δ* represents the deviation from the nominal values. The initial speed of the response is given by the pre-factor of the *δx* term. This is larger in the negative feedback circuit than in the no feedback circuit. Even in the negative feedback circuit, the initial speed is larger with stronger feedback (smaller *K*) than with weaker feedback (larger *K*). Therefore, the initial difference can be larger even if the overall response is more contracting.

## III. Conclusion

We showed that temperature robustness can be facilitated by the structure of a biomolecular circuit. Using a benchmark negative feedback circuit, we presented three main results. First, we found that the difference in the response to a step change in temperature in a mathematical model of the negative feedback circuit was smaller than in a circuit without feedback. Second, we experimentally verified this prediction by synthesizing circuits with and without feedback. Third, we discussed a counterintuitive finding that even though the negative feedback circuit is more robust than the no feedback circuit, the initial rate of difference in response is actually larger in the negative feedback circuit.

Our results show how temperature robustness can be implemented through the circuit structure. Mechanisms for designing circuit structure are relatively simpler than tuning the temperature dependence of the circuit parameters. Therefore, this principle should be useful in designing temperature robust biomolecular circuits as well as in understanding temperature robustness in naturally occurring biomolecular circuits.

## IV. Materials and methods

The construction of the plasmids and strains as well as the workflow for experimental measurements and data analysis is summarized below.

### A. Plasmids and Strains

We used DNA synthesis to construct the circuits with negative feedback and without the feedback. The circuit and the Plasmid map are available in Supplementary Figure 3. For the negative feedback construct, we synthesized the fragment *P*_*tet*_-*TetR* :: *sf GFP* -*rrnB*1*B*2*T* 1 containing the *P*_*tet*_ promoter, a fusion of the *TetR* protein and the Green Fluorescent Protein *sf GFP* , and a terminator is used from [28]. For the construct without feedback, we synthesized the fragment *P*_*tet*_-*sf GFP* -*rrnB*1*B*2*T* 1. Both fragments were inserted into a vector based on *pUC*57 using the restriction sites *EcoRI* and *SalI*. The vector had a *pUC*57 backbone with ampicillin resistance and had been modified so that there was another terminator *TBS*7 upstream of the insert. The Plasmid description and sequences are available in the Table 1 and Table 2 of the Supplementary Material, respectively. Both plasmids were transformed into the *E*.*coli* strain *DH*5*α*.

### B. Measurements

For experimental measurements, an overnight culture was prepared by inoculating 1 *mL* Luria-Bertani (LB) medium supplemented with 1 *µL* of 200 *µg/mL* ampicillin with a bacterial colony. This was incubated for 16 hours at 29°C with shaking. The overnight cultures were subsequently diluted 1 : 100 in fresh LB medium containing ampicillin and grown at 29°C with orbital shaking for 2 hours, respectively. The inducer Anhydrotetracycline (*aTc*) was added at different concentrations of 0, 15, 25, 50*ng/mL* to separate aliquots of diluted bacterial culture. The samples were transferred to individual wells of 96-well microplate (Thermo Scientific), each in triplicate. The plate was incubated in a microplate reader (BioTek Synergy H1) equipped with double orbital shaking (2 minute interval) at 29°C. The temperature step was given by maintaining cultures at 29°C for the initial 2 hours, followed by a step increase to 37°C for the remaining 12 hours of measurement. Measurements of Optical Density (600*nm*) and Fluorescence (excitation: 485*nm*; emission: 520*nm*) were recorded at 5-minute intervals throughout the experiment. For experiments in minimal medium, the overnight cultures were diluted into the minimal medium (1 × *M* 9 Salts, 1.0% Glucose, 0.1% Casamino Acids, 0.5*µg/mL* Thiamine, 0.2*mM* Magnesium Sulfate, 0.1*mM* Calcium Chloride, adjusted to pH 7.0 from Teknova) containing ampicillin. The experimental measurements were repeated thrice.

### C. Data Analysis

The data were analysed in MATLAB. The mean of the background media used is subtracted from these raw optical density and fluorescence data sample means for the per-cell measurements.

## Supplementary Materials

The supplementary material for this article is available at this link.

## Acknowledgement

The authors thank Mr Arun Thapa, Ms Ngangom Pravina Devi, and Ms Sakshi (PhD scholars of the Department of Biochemical Engineering and Biotechnology, IIT Delhi) for their help in providing access to some of the laboratory equipment of their lab.

