## Supplementary Figure for "A Structural Design Principle for Temperature Robustness in Biomolecular Circuits"

### Contents

A. Supplementary Experimental Data (Supplementary Figures 1-2) 3

B. Circuits and Plasmid Maps (Supplementary Figure 3) 5

C. Plasmids description 6

D. Sequence Information 6

#### A. Supplementary Experimental Data

##### Supplementary Figures 1

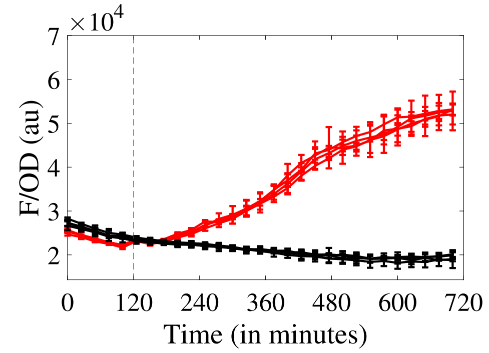


(a)

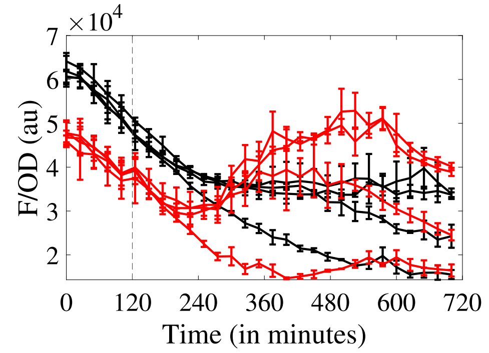


(b)

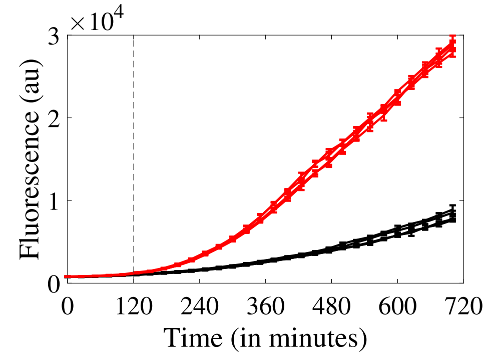


(c)

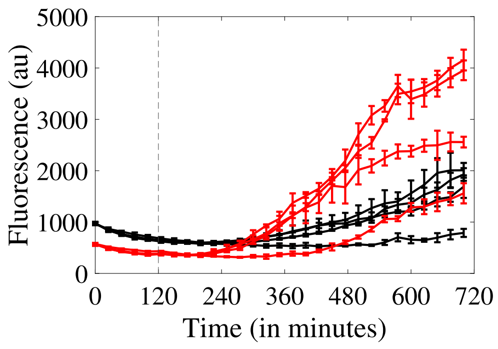


(d)

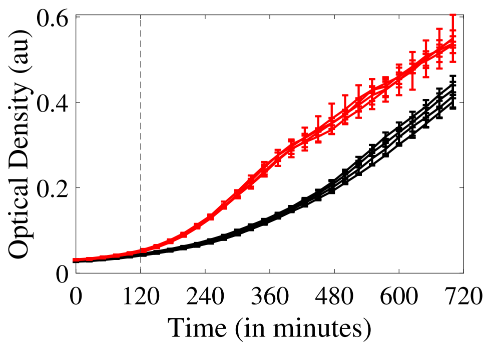


(e)

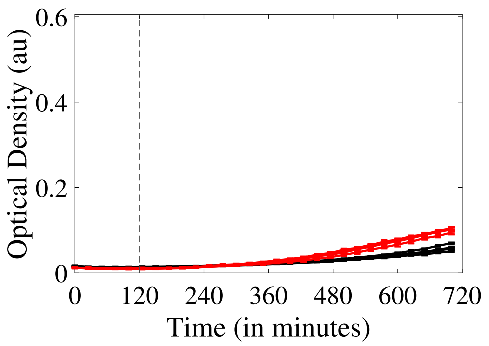


(f)

Figure 1: Experimental measurements of the temperature step response (red curves) and control response (black curves) in Minimal media. (a) and (b) are the responses of no and negative feedback, respectively. (c) and (d) are the fluorescence measurements of the response in panel a and b, (e) and (f) are the optical density measurements of the response in panel a and b. These curves are with different aTc concentrations.

**Supplementary Figures 2**

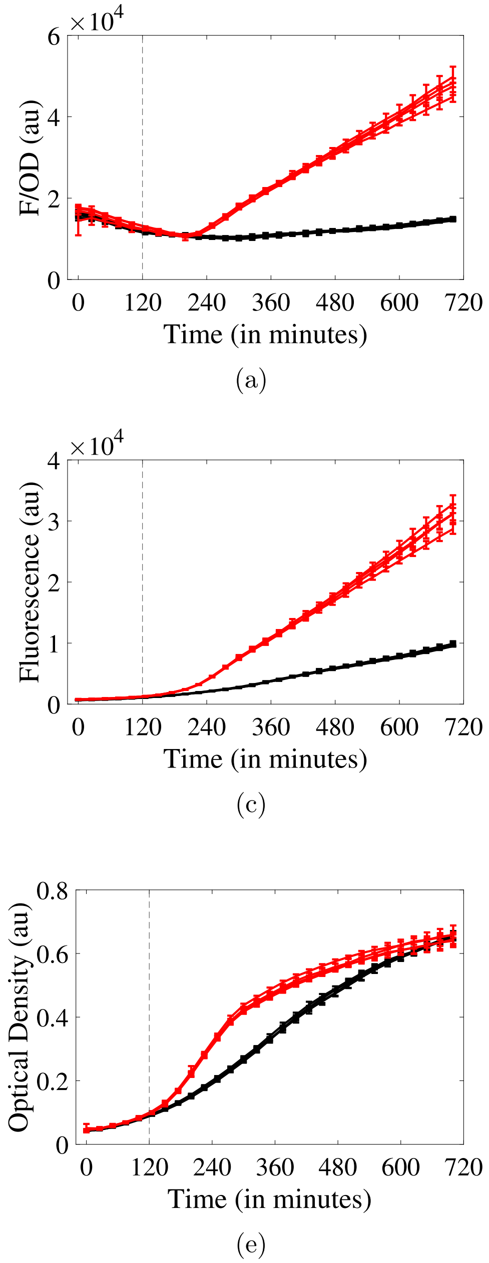


Figure 2: Experimental measurements of the temperature step response (red curves) and control response (black curves) in LB media. (a) and (b) are the responses of no and negative feedback, respectively. (c) and (d) are the fluorescence measurements of the response in panel a and b, (e) and (f) are the optical density measurements of the response in panel a and b. These curves are with different aTc concentrations.

#### B. Circuits and Plasmid Maps

##### Supplementary Figure 3

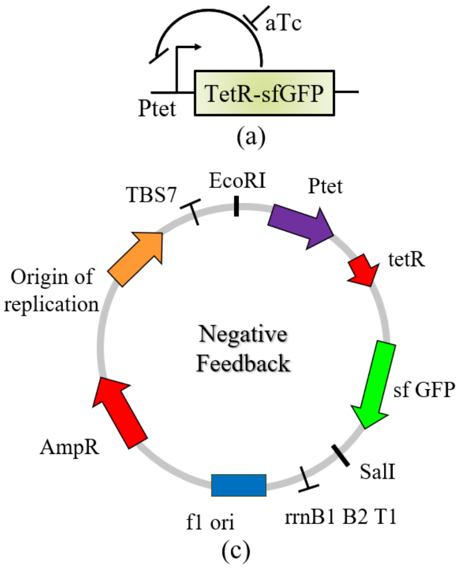


Figure 3: The circuits and their plasmid schematic with necessary components used in this experimental study.

#### C. Plasmids description

Table 1: Plasmids description

| S. No | Circuit Description | Antibiotic |
|---|---|---|
| 1. | $P_{tet}$-$TetR :: sfGFP$-$rrnB1B2T1$ | Ampicillin |
| 2. | $P_{tet}$-$sfGFP$-$rrnB1B2T1$ | Ampicillin |

#### D. Sequence Information

Table 2: Sequences

| Component | Sequences | Reference |
|---|---|---|
| $P_{tet}$ | TCCCTATCAGTGATAGAGATTGACATCCCTATCAGTGATAGAGATACTGAGCAC | 1 |
| $TetR - sfGFP$ | ATGTCCAGATTAGATAAAAGTAAAGTGATTAACAGCGCATTAGAGCTGC<br>TTAATGAGGTCGGAATCGAAGGTTTAACAACCCGTAAACTCGCCCAGAA<br>GCTAGGTGTAGAGCAGCCTACATTGTATTGGCATGTAAAAAATAAGCGG<br>GCTTTGCTCGACGCCTTAGCCATTGAGATGTTAGATAGGCACCATACTC<br>ACTTTTGCCCTTTAGAAGGGGAAAGCTGGCAAGATTTTTTACGTAATAA<br>CGCTAAAAGTTTTAGATGTGCTTTACTAAGTCATCGCGATGGAGCAAAA<br>GTACATTTAGGTACACGGCCTACAGAAAAACAGTATGAAACTCTCGAAA<br>ATCAATTAGCCTTTTTATGCCAACAAGGTTTTTCACTAGAGAATGCATT<br>ATATGCACTCAGCGCTGTGGGGCATTTTACTTTAGGTTGCGTATTGGAA<br>GATCAAGAGCATCAAGTCGCTAAAGAAGAAAGGGAAACACCTACTACTG<br>ATAGTATGCCGCCATTATTACGACAAGCTATCGAATTATTTGATCACCA<br>AGGTGCAGAGCCAGCCTTCTTATTCGGCCTTGAATTGATCATATGCGGA<br>TTAGAAAAACAACTTAAATGTGAAAGTGGGTCCCGTAAAGGCGAAGAGC<br>TGTTCACTGGTGTCGTCCCTATTCTGGTGGAACTGGATGGTGATGTCAA<br>CGGTCATAAGTTTTCCGTGCGTGGCGAGGGTGAAGGTGACGCAACTAAT<br>GGTAAACTGACGCTGAAGTTCATCTGTACTACTGGTAAACTGCCGGTAC<br>CTTGGCCGACTCTGGTAACGACGCTGACTTATGGTGTTCAGTGCTTTGC<br>TCGTTATCCGGACCATATGAAGCAGCATGACTTCTTCAAGTCCGCCATG<br>CCGGAAGGCTATGTGCAGGAACGCACGATTTCCTTTAAGGATGACGGCA<br>CGTACAAAACGCGTGCGGAAGTGAAATTTGAAGGCGATACCCTGGTAAA<br>CCGCATTGAGCTGAAAGGCATTGACTTTAAAGAAGACGGCAATATCGAT<br>ACTGGGCCATAAGCTGGAATACAATTTTAACAGCCACAATGTTTACATC<br>ACCGCCAACAAAAAAATGGCATTAAAGCGAATTTTAAAATTCGCCACAA<br>CGTGGAGGATGGCAGCGTGCAGCTGGCTGATCACTACCAGCAAAACACT<br>CCAATCGGTGATGGTCCTGTTCTGCTGCCAGACAATCACTATCTGAGCA<br>CGCAAAGCGTTCTGTCTAAAGATCCGAACGAGAAACGCGATCATATGGT<br>TCTGCTGGAGTTCGTAACCGCAGCGGGCATCACGCATGGTATGGATGAA<br>CTGTACAAAGGCTCCGGCTCCGGCTCCCACCATCACCATCACCATTGA | 1 |
| $sfGFP$ | ATGCGTAAAGGCGAAGAGCTGTTCACTGGTGTCGTCCCTATTCTGGTGGAACT<br>GGATGGTGATGTCAACGGTCATAAGTTTTCCGTGCGTGGCGAGGGTGAAGGTG<br>ACGCAACTAATGGTAAACTGACGCTGAAGTTCATCTGTACTACTGGTAAACTG<br>CCGGTACCTTGGCCGACTCTGGTAACGACGCTGACTTATGGTGTTCAGTGCTT<br>TGCTCGTTATCCGGACCATATGAAGCAGCATGACTTCTTCAAGTCCGCCATGC<br>CGGAAGGCTATGTGCAGGAACGCACGATTTCCTTTAAGGATGACGGCACGTAC<br>AAAACGCGTGCGGAAGTGAAATTTGAAGGCGATACCCTGGTAAACCGCATTGA<br>GCTGAAAGGCATTGACTTTAAAGAAGACGGCAATATCCTGGGCCATAAGCTGG<br>AATACAATTTTAACAGCCACAATGTTTACATCACCGCCGATAAACAAAAAAAT<br>GGCATTAAAGCGAATTTTAAAATTCGCCACAACGTGGAGGATGGCAGCGTGCA<br>GCTGGCTGATCACTACCAGCAAAACACTCCAATCGGTGATGGTCCTGTTCTGC<br>TGCCAGACAATCACTATCTGAGCACGCAAAGCGTTCTGTCTAAAGATCCGAAC<br>GAGAAACGCGATCATATGGTTCTGCTGGAGTTCGTAACCGCAGCGGGCATCAC<br>GCATGGTATGGATGAACTGTACAAA | 1 |
| $rrnB1B2T1$ term terminator | GGCTCCGGCTCCGGCTCCCACCATCACCATCACCATTGATAACTCGAGCCCCA<br>AGGGCGACACCCCATAATTAGCCCGGGCGAAAGGCCCAGTCTTTCGACTGAGC<br>CTTTCGTTTTATTTGATGCCTGGCAGTTCCCTACTCTCGCATGGGGAGTCCCC<br>ACACTACCATCGGCGCTACGGCGTTTCACTTCTGAGTTCGGCATGGGGTCAGG<br>TGGGACCACCGCGCTACTGCCGCCAGGCAAA | 1 |
